# Reward processing in children with Affective Dysregulation

**DOI:** 10.1101/2024.10.02.616272

**Authors:** Pascal-M Aggensteiner, Francisca Giller, Nathalie Holz, Anna Kaiser, Christine Igel, Sarah Hohmann, Manfred Döpfner, Veit Roessner, Christian Beste, Tobias Banaschewski, Daniel Brandeis, the ADOPT Consortium

## Abstract

**Introduction:** Affective dysregulation (AD) in children is characterized by irritability, anger, and frequent intense temper outbursts. Considerable evidence implies altered processing of frustration about missed rewards, but few studies investigated the preceding and thus potentially predictive reward anticipation and initial delivery processing in children with AD.

**Methods:** A total of 103 children aged 8 to 12 years (50 with AD and 53 without AD) were examined during a monetary reward anticipation task with event-related potential (ERP) components resolving reward anticipation (cue-CNV [Contingent Negative Variation]) and reward delivery phases (Reward Positivity and Feedback-Related Negativity). All components were analyzed by repeated measures ANOVA. Regression analyses also evaluated the associations between those ERP components and dimensional AD symptoms.

**Results:** Children with AD showed attenuated anticipatory reward processing compared to No-ADs. The CNV at fronto-central site (FCz) showed a significant group effect (No-AD>AD, p=0.017). Post-hoc test showed that this group difference was stronger for the cue monetary condition (monetary cue: p=.007, d=0.56, verbal cue: p=.901, d=0.16), and that only the No-AD group showed a significant difference between conditions (p<0.001). No significant effects were obtained for the delivery phase. Regression analysis showed that a reduced anticipatory CNV at FCz significantly explained AD symptoms, and that anger/irritability and anxiety/depressive symptoms predicted a reduced anticipatory CNV at FCz.

**Conclusion:** This neurophysiological characterization of reward anticipation and delivery in children with AD demonstrates altered neural activity in AD during anticipation of reward rather than following the delivery (or omission) of the reward itself. Our results highlight that altered reward anticipation in AD can occur outside frustration-prone tasks or settings, and underline the important role of both anger/irritability and anxiety/depressive symptoms in the pathophysiology of AD for atypical reward anticipation.

## 1. Introduction

Anger, aggression and irritability are key elements of affective dysregulation (AD) in children. AD is characterized by a general irritable mood plus age-and-situation-inappropriate temperament outbursts with aggressive tendencies (Brotman et al., 2017). The terms irritability and AD which are often used synonymously (Döpfner et al., 2019a; Waltereit et al., 2019), are increasingly accepted as a transdiagnostic concept bridging several mental disorders including disruptive mood dysregulation disorder (DMDD), oppositional defiant disorder (ODD), conduct disorder (CD) and attention deficit hyperactive disorder (ADHD) (Jensen *et al*. 2007; Rich *et al*., 2011; Burke *et al*., 2014; Shaw *et al*., 2014; Ambrosini *et al.,* 2013).

Altered reward processing has been proposed as a key mechanism of irritability (Brotman et al., 2017) or AD (Döpfner et al., 2019b). Although changes in reward processing with AD have been described (Avenevoli et al., 2015; Kessel et al., 2016; Leibenluft & Stoddard, 2013), this research mainly focused on the delivery phase during frustrating situations where expected rewards are not received, aligning with the frustrative non-reward, rather than the reward anticipation construct also listed in the Research Domain Criteria (RDoC) (Insel et al., 2010; Musser & Raiker, 2019). Several EEG studies, although mainly in healthy controls or community samples, demonstrated significant electrophysiological alterations with AD-related symptoms during the absence of an expected reward (frustrative non-reward, i.e. Deveney et al., 2019; Waxmonsky et al., 2019) and the receipt of a rewarding stimulus (reward sensitivity, i.e. [Kessel et al., 2016; Waxmonsky et al., 2022]). However, the findings regarding the reward delivery phase remain inconsistent, indicating either hypo- or hyperactivity, which is likely partially attributed to the heterogeneous methodology (Hennefield et al., 2022) and definitions of AD-symptoms across studies.

Surprisingly, no ERP studies have yet assessed whether AD in groups meeting stringent criteria is associated with reward anticipation preceding reward attainment or frustration, although reward anticipation is a core aspect of reward processing, an established RDoC construct, and known to partially predict reward processes in healthy controls (Zubovics et al., 2020). Our study therefore, specifically investigates anticipatory reward processes in well-characterized young children with AD.

Reward anticipation is typically measured in reward processing tasks by the CNV, a slow cortical potential linked to anticipation, expectation and cognitive preparation in a wide range of tasks. This CNV builds up approximately one second after a reward cue until the reward target appears, precedes feedback, and is stronger (more negative) in response to the anticipation of more salient rewards (Boecker et al., 2014; Broyd et al., 2012). It is probably generated in the dorsal anterior cingulate, frontal cortex and midbrain dopaminergic nuclei (Albrecht et al., 2013, Plichta et al., 2013), susceptible to dopaminergic modulation (Linssen et al., 2011). Studies assessing reward anticipation reported that participants self-reporting more difficulties in regulating emotions showed weaker anticipatory ERPs during a monetary incentive delay task (Zubovics et al., 2021) in high school students and in adults with depressive symptoms (Ait Oumeziane et al., 2019).

Regarding the components related to reward delivery, the Reward Positivity (RewP) and Feedback-Related Negativity (FRN) are components which typically show stronger activity to successful trials (Foti et al., 2011; Pfabigan et al., 2014), and have been associated with increased ventral striatum and medial prefrontal cortex brain activity (Carlson et al., 2011). Neurophysiological deviations in reward delivery are somewhat better investigated in irritability or AD. Preschoolers diagnosed with DMDD were assessed later in preadolescence during a monetary reward task. This study found an association between DMDD symptoms assessed at age 3 and enhanced RewP to monetary rewards (Kessel et al., 2016), consistent with previous work on irritability in middle-late childhood (Dougherty et al. 2015). In young adults, higher irritability was associated with reduced response to loss feedback, but across the whole sample, trait irritability was unrelated to reward responsivity (Deveney, 2019). An association between CD and reward responsivity was found to be moderated by irritability, such that youth with high CD symptom had less positive reward responsivity to win trials and more positive responsivity in loss trials, but this pattern was only observed in youth with high irritability scores (Waxmonsky et al., 2022). Importantly, none of the above mentioned studies was based on a case-control comparison and only the study of Waxmonsky et al., 2022 included a clinical sample, highlighting the need to follow this up in a larger cohort comparing those ERPs between AD and No-AD peers.

Our study aims to contribute to increase our understanding of the relationship between reward processing and AD considering both the anticipatory CNV and the delivery RewP/FRN. We hypothesize that AD will show reduced anticipatory CNV and RewP/FRN activity, compared to a control group, reflecting altered cognitive anticipation and reward processing. Additionally, by adopting a dimensional approach that emphasizes measurable and continuous neurophysiological markers, we aim to assess how variations in reward-related activity may explain AD symptoms, and by identifying the underlying neural mechanisms, we may enhance diagnostics and deepen our comprehension of the pathophysiology of AD. This potentially offers enhanced diagnostic precision and more targeted therapeutic interventions and increases our understanding of the phenotypic representation and the associated pathophysiology related to the conceptualization of AD.

## 2. Methods

### 2.1. Study design and participants

Participants were part of the randomized, controlled trial ADOPT (Affective Dysregulation—Optimizing Prevention and Treatment) (Döpfner et al., 2019a). The participants were randomly recruited through the residents’ registration offices of four cities. Following screening for AD, children were classified into two groups: high AD (top 10% of raw scores) and low AD (bottom 10%). We selected this 10% cut-off to approximate the AD prevalence rates of up to 9% reported in epidemiological studies. Families of all children identified with AD, along with a randomly selected sample of families of children without AD, were invited to participate in a comprehensive assessment at one of five study centers. All families with children meeting the additional inclusion criteria were invited to take part in the subsequent treatment study, where they received either an AD-specific treatment or standard care. Meanwhile, families without AD were monitored as a comparison group (ADOPT study, Döpfner et al., 2019). Here we report the data of the subgroup that participated in the ADOPT-Neurobiology study, which was performed at the Technische Universität Dresden and the Central Institute of Mental Health, Mannheim. A total of 138 children (aged 8-12) participated in this EEG assessment. The full assessment battery consisted of three tasks: Emotional Stroop (Giller et al., 2021), Reward Anticipation (Kirsch et al., 2003) and Affective Posner Cueing Task (Rich et al., 2007) in addition to two 3 min resting state measurements (eyes open and eyes closed). Ethical approval for the study was obtained from the local ethics committee. Written consent was obtained from the children and their legal guardians.

Children with an IQ of less than 80, with current or past traumatic brain injuries, with neurological diseases and/or with a confirmed diagnosis of bipolar disorder, manic episode, severe depression, or autism spectrum disorder were not included in the study. Moreover, affective dysregulation should not be better explained by other disorders such as obsessive-compulsive disorder, posttraumatic stress disorder, persistent depressive disorder, or substance use disorders. Assignment in the clinical participant group was determined by a certified clinician informed by the Diagnostic Tool for Affective Dysregulation in Children (DADYS, Treier et al., 2023; Junghänel et al., 2022) as evaluated by a parent/guardian and by a clinician based on a parent interview. Comorbidities such as ADHD, ODD, or CD or stimulant medication (e.g. methylphenidate) were not a reason for exclusion and are shown in Table 1. Children who were currently undergoing or planned to undergo intensive behavioral therapy were not included in the study. All children had a vision that was normal or corrected to normal.

**Table 1.**
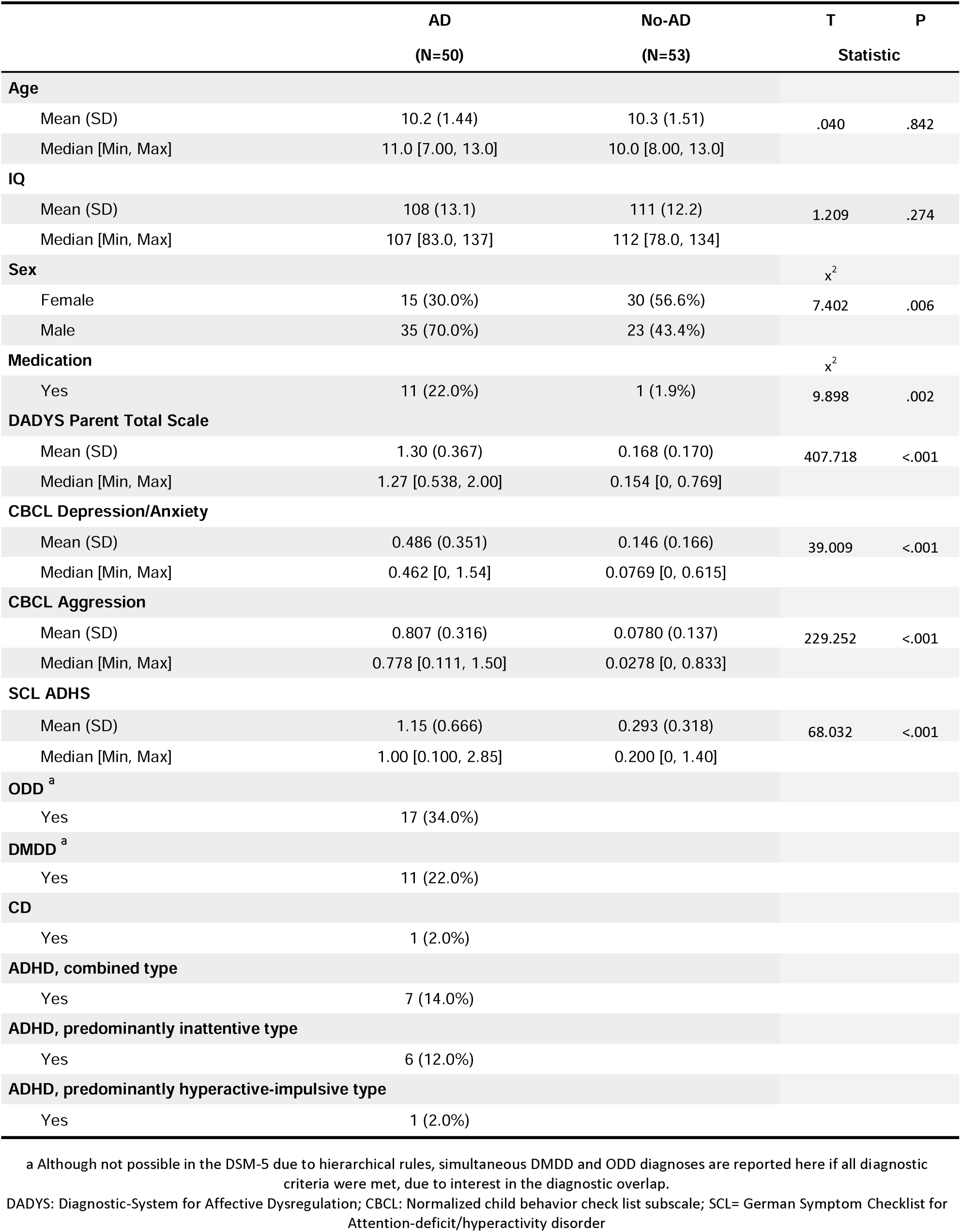
Sample characteristics.

|  | AD<br>(N=50) | No-AD<br>(N=53) | T<br>Statistic | P |
| --- | --- | --- | --- | --- |
| **Age** |  |  |  |  |
| Mean (SD) | 10.2 (1.44) | 10.3 (1.51) | .040 | .842 |
| Median [Min, Max] | 11.0 [7.00, 13.0] | 10.0 [8.00, 13.0] |  |  |
| **IQ** |  |  |  |  |
| Mean (SD) | 108 (13.1) | 111 (12.2) | 1.209 | .274 |
| Median [Min, Max] | 107 [83.0, 137] | 112 [78.0, 134] |  |  |
| **Sex** |  |  |  |  |
| | | | $\chi^2$ | |
| Female | 15 (30.0%) | 30 (56.6%) | 7.402 | .006 |
| Male | 35 (70.0%) | 23 (43.4%) |  |  |
| **Medication** |  |  |  |  |
| | | | $\chi^2$ | |
| Yes | 11 (22.0%) | 1 (1.9%) | 9.898 | .002 |
| **DADYS Parent Total Scale** |  |  |  |  |
| Mean (SD) | 1.30 (0.367) | 0.168 (0.170) | 407.718 | <.001 |
| Median [Min, Max] | 1.27 [0.538, 2.00] | 0.154 [0, 0.769] |  |  |
| **CBCL Depression/Anxiety** |  |  |  |  |
| Mean (SD) | 0.486 (0.351) | 0.146 (0.166) | 39.009 | <.001 |
| Median [Min, Max] | 0.462 [0, 1.54] | 0.0769 [0, 0.615] |  |  |
| **CBCL Aggression** |  |  |  |  |
| Mean (SD) | 0.807 (0.316) | 0.0780 (0.137) | 229.252 | <.001 |
| Median [Min, Max] | 0.778 [0.111, 1.50] | 0.0278 [0, 0.833] |  |  |
| **SCL ADHS** |  |  |  |  |
| Mean (SD) | 1.15 (0.666) | 0.293 (0.318) | 68.032 | <.001 |
| Median [Min, Max] | 1.00 [0.100, 2.85] | 0.200 [0, 1.40] |  |  |
| **ODD<sup>a</sup>** |  |  |  |  |
| Yes | 17 (34.0%) |  |  |  |
| **DMDD<sup>a</sup>** |  |  |  |  |
| Yes | 11 (22.0%) |  |  |  |
| **CD** |  |  |  |  |
| Yes | 1 (2.0%) |  |  |  |
| **ADHD, combined type** |  |  |  |  |
| Yes | 7 (14.0%) |  |  |  |
| **ADHD, predominantly inattentive type** |  |  |  |  |
| Yes | 6 (12.0%) |  |  |  |
| **ADHD, predominantly hyperactive-impulsive type** |  |  |  |  |
| Yes | 1 (2.0%) |  |  |  |

a Although not possible in the DSM-5 due to hierarchical rules, simultaneous DMDD and ODD diagnoses are reported here if all diagnostic criteria were met, due to interest in the diagnostic overlap.
DADYS: Diagnostic-System for Affective Dysregulation; CBCL: Normalized child behavior check list subscale; SCL= German Symptom Checklist for Attention-deficit/hyperactivity disorder

### 1.1. Reward anticipation task

Adapted from previous versions of the MID task (Kirsch et al., 2003), the paradigm was designed to measure reward anticipation and delivery of a monetary reward, and has been reported to activate the ventral striatum reliably and robustly (Boecker et al., 2014; Plichta et al., 2013). The visual information of the paradigm was presented on a monitor using Presentation software (Neurobehavioral Systems). As depicted in the supplementary figure 1, the task requires the participant to respond by a fast button press once presented with a flash target. A fast press secures the target-indicated reward. The smiley target cue indicates a potential monetary reward (0.50 Euros) and the scrambled smiley cue, a verbal feedback (i.e. “Fast reaction!”), which served as the control condition. To encourage further engagement with the task, boost monetary trials of 2 Euros were included circa every eighth win trial. Participants were reminded of their overall account balance after every trial. The paradigm presented a total of 50 trials for each condition in a pseudo-randomized order. Jittering of the cue duration was between 3-5s. The allocated reaction time window for a successful trial was adapted to account for between-subject differences (max. 1 s), intended to support a comparable number of successful trials between participants (60% of trials). This time window remained constant between both conditions. Following measurement sessions, participants were paid their final earned balances (for details, see supplementary Figure 1).

## 3. Assessment of Demographic Information and Clinical Characterization

Demographic information was assessed before the EEG assessment. IQ was measured via a 4 subtests of German WISC IV (Petermann & Petermann, 2007), namely Symbol Search and Vocabulary along with Matrix Reasoning and Block Design.

### DADYS Questionnaire

The DADYS (Diagnostikum für Affektive Dysregulation [Diagnostic-System for Affective Dysregulation], Görtz-Dorten & Döpfner, 2021), is a newly developed diagnostic system for the multimodal assessment of affective dysregulation in children. Our analysis is based on the validated parent questionnaire (DADYS-PQ, 36 items, Junghänel, Thöne, et al., 2022). This questionnaire contains an array of items based on AD-associated symptoms—functional impairment, irritability/emotional impulsivity, anger/irritability, positive emotionality and exuberance. Items of all DADYS instruments are rated on a 4-point Likert scale (0=not present to 3=very strong).

### CBCL Questionnaire

Using the Child Behavior Checklist 6/18 (CBCL; Achenbach, 2001), parents were able to report on the problems with behavior and emotions of their child. This analysis used the scales from this questionnaire, which measure anxiety/depression, attention difficulties and aggressive behavior (Achenbach & Rescorla, 2014, Döpfner et al., 2014).

### SCL-ADHD Questionnaire

Parent reports of ADHD symptoms were collected via the SCL-ADHS form of the German Symptom Checklist for Attention-deficit/hyperactivity disorder from the German diagnostic system for mental disorders in children and adolescents based on the ICD-10 and DSM-5 (DISYPS-III, Döpner et al., 2017). The questionnaire comprised 20 items, which are rated on a four-point likert scale ranging from 0 (not at all) to 3 (very much).

### 1.2. EEG data acquisition and preprocessing

Participants were seated in a comfortable chair and fitted with a 60-channel EEG cap (Brain Products Inc.). The EEG was recorded with a sampling rate of 500 Hz using a 72-channel BrainAmp amplifier (Brain Products Inc.) and BrainVision Recorder software (Brain Products Inc.). An equidistant 60 Ag–AgCl-EEG setup was used with the ground electrode at θ = 58, φ = 78 (between AFz and AF2) and the recording reference electrode Cz. Acquisition started after impedances for all channels were reduced to below 50 kΩ following standard data collection procedures (Ferree et al., 2001; Kappenman & Luck, 2010). One electrode was placed below the outer canthus of each eye (horizontal electrooculogram HEOG I & II) and one below the left eye (vertical electrooculogram VEOG I).

Offline processing was performed with Brain Vision Analyzer 2.2.0 (BrainProducts, Gilching Germany). The continuous EEG was filtered with 0.1–30 Hz, 24 db/oct Butterworth filters, broad artefacts were eliminated after visual inspection and heavily affected channels were interpolated using the spline type. Ocular artefacts were removed by an independent component analysis (ICA, infomax algorithm). Data were re-referenced to the average and subsequently checked for remaining artefacts. Automated artifact rejection was applied for all segments using amplitude differences above 200 μV in a 200 ms time interval as well as activity ± 150 μV as rejection criteria. For the anticipation phase, the CNV was the primary ERP measured from 2000-3000s at Cz and FCz, which commonly show the highest (more negative) amplitude in this task (Boecker et al., 2014; Plichta et al., 2013) and are typically more frontal at younger ages. Based on the visual inspection of scalp grand-averaged topographic plots and ERP waveforms, which are commonly used to study RewP/FRN (Hennefield et al., 2022), the RewP was calculated between 175-275ms and the FRN between 250-350ms, and the mean amplitude was extracted for selected electrodes (RewP: AFz, AF3, Fz, and AF4; FRN: FCz, Cz CPz, CP1 and CP2). The difference in activity between win and no win was calculated for each condition (verbal and monetary).

### 3.1. Statistical analysis

Sample characteristics were analyzed using analysis of variance (ANOVA) and chi-square tests, when appropriate. Behavioral data (Reaction time) were analyzed using repeated measures ANOVA, with one within-factor for condition (monetary and verbal) and one between-factor for group (AD and No-AD). ERP reward anticipation data were analyzed using repeated measures ANOVA, with two within-factors for condition (monetary and verbal) and electrodes (Cz and FCz) and one between-factor for group (AD and No-AD). EEG reward delivery data (RewP and FRN separately) were analyzed using repeated measures ANOVA, with one within-factor for condition (monetary and verbal) and one between-factor for group (AD and No-AD). All analyses were adjusted for sex as a covariate of non-interest. Additionally, a sensitivity analysis was performed, including site and medication as covariates, and all analyses were repeated using only participants with available behavioral data. Coheńs *d* and partial η2 effect sizes were reported (Cohen, 1988)

To determine the extent to which EEG reward-related activity can explain AD symptoms, two stepwise linear regression models were applied including both groups. The first regression assessed how much variance can be explained by EEG activity, with sex and medication added as control variables in the first step. In the second step, EEG activity for reward anticipation at Cz and FCz was added, and in the third step, the RewP for the reward delivery phase were introduced. The same model was repeated, adding ADHD, aggression and anxiety/depressive symptoms as a second step. In the third step, reward anticipation at Cz and FCz was included, and in the fourth step, the RewP or FRN for the reward delivery phase was introduced.

Subsequently, we aimed to assess which symptom dimensions of affective dysregulation or its comorbid symptoms can predict the reward EEG activity in separate regression models. These models were also controlled for sex and medication, with the dependent variables being CNV reward anticipation activity at FCz, RewP or FRN, and the independent variables being key symptoms of affective dysregulation or comorbidities, such as aggression, anxiety/depression, and ADHD. Correction for multiple testing was performed for these regression analyses applying a Bonferroni correction for 8 test (0.05/8=0.00625). All analysis were performed with SPSS, Version 26 and R Studio v4.1.2.

## 4. Results

### 4.1. Sample characteristics

Table 1 shows the sample characteristics. From the 138 participants available for EEG analysis, 35 participants could not be included in the final analysis because of errors in recordings or poor quality of the EEG data (less than 20 segments/epochs for each condition). For analysis of the reward anticipation phase, 103 participants (53 No-AD and 50 AD) were included. Compared to No-AD, the AD group consisted of more males (x^2^=7.402, p < 0.001). For the reward delivery phase, slightly fewer participants (RewP, n= 96, No-AD= 52, AD=44; FRN, n=101, No-AD=53, AD=48) could be analyzed due to higher presence of movement artifacts and the presence of one outlier. For the reaction time data 93 participants could be analyzed (No-AD=52, AD=41) due to further technical problems. Furthermore, no significant differences were found for the baseline characteristics and symptom severity of the excluded participant’s vs the included ones.

### 4.2. Behavioral data

Reaction Times (RT) were faster following monetary cues [F(1,90)=7.012, p=0.010, part. η2=0.072]. Both groups performed similarly and no significant interaction and group effect was found (all p>0.489). For details, see supplementary Figure 2.

### 4.3. Anticipation

A significant group x condition interaction [F(1,100)=5.409, p=0.022, part. η2=0.051] revealed that the AD group had reduced anticipatory CNV amplitude. Post-hoc test showed that this between-group difference was stronger for the cue monetary condition (monetary cue: p=.007, d=0.56, verbal cue: p=.901, d=0.16), and that only the No-AD group showed a significant difference between conditions (p<0.001)]. A significant group x channel interaction [F(1,100)=4.052, p=0.047, part. η2=0.039] revealed that the group differences were more pronounced at the fronto-central electrode (FCz, p=.017). No other significant effects or interactions emerged.

For details see Figure 1 and Table 2

**Figure 1.**
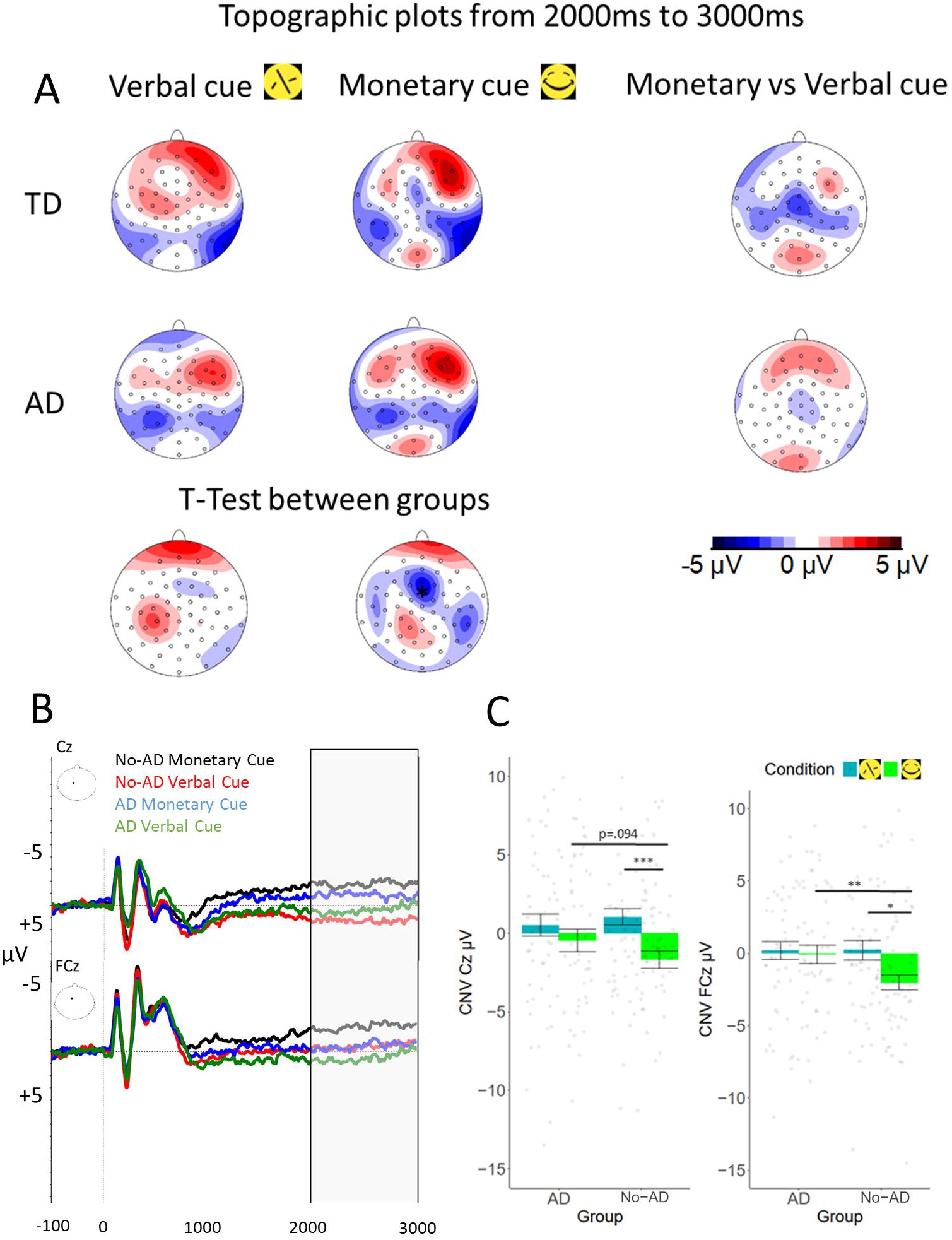
Reward anticipation CNV 2000ms – 3000ms

**Table 2.** Descriptive statistics of ERPs.

|  | AD<br>(N=50) | No-AD<br>(N=53) | Cohen's *d* |
| --- | --- | --- | --- |
| **Monetary cue CNV (FCz)** |  |  |  |
| Mean (SD) | 0.469 (5.13) | -2.02 (3.78) | .56** |
| Median [Min, Max] | -0.404 [-8.83, 16.3] | -2.32 [-13.6, 7.41] | 95% CI [0.16, 0.95] |
| **Verbal cue CNV (FCz)** |  |  |  |
| Mean (SD) | 0.784 (5.11) | -0.0770 (5.35) | .16 |
| Median [Min, Max] | 0.489 [-11.3, 18.4] | -0.444 [-16.1, 9.88] | 95% CI [-0.22, 0.55] |
| **Monetary cue CNV (Cz)** |  |  |  |
| Mean (SD) | -0.233 (5.22) | -1.98 (4.53) | .36 |
| Median [Min, Max] | -0.665 [-12.0, 10.8] | -1.68 [-17.9, 6.55] | 95% CI [-0.03, 0.75] |
| **Verbal cue CNV (Cz)** |  |  |  |
| Mean (SD) | 0.513 (4.97) | 1.23 (3.91) | -.16 |
| Median [Min, Max] | 1.10 [-13.5, 9.15] | 0.166 [-5.15, 11.0] | 95% CI [-0.55, 0.23] |
| **Verbal RewP** |  |  |  |
| Mean (SD) | 1.22 (3.86) | 1.73 (5.33) | -.14 |
| Median [Min, Max] | 0.830 [-5.56, 12.6] | 1.83 [-11.8, 15.0] | 95% CI [-0.51, 0.29] |
| Missing | 6 (12.0%) | 1 (1.9%) |  |
| **Monetary RewP** |  |  |  |
| Mean (SD) | -0.214 (3.50) | 1.65 (4.59) | -.45° |
| Median [Min, Max] | 0.0465 [-9.08, 8.02] | 0.897 [-7.02, 15.4] | 95% CI [-0.86, -0.05] |
| Missing | 6 (12.0%) | 1 (1.9%) |  |
| **Verbal FRN** |  |  |  |
| Mean (SD) | 1.44 (3.12) | 0.673 (2.89) | .25 |
| Median [Min, Max] | 1.27 [-4.28, 7.97] | 0.542 [-4.59, 7.16] | 95% CI [-0.14, 0.65] |
| Missing | 2 (4.0%) | 0 (0%) |  |
| **Monetary FRN** |  |  |  |
| Mean (SD) | 0.0715 (3.18) | 0.300 (2.71) | -.08 |
| Median [Min, Max] | -0.336 [-7.59, 6.19] | 0.112 [-5.36, 5.73] | 95% CI [-0.47, 0.31] |
| Missing | 2 (4.0%) | 0 (0%) |  |

AD: Affective dysregulation, CNV: Contingent negative variation, RewP: Reward positivity,
FRN: Feedback related negativity, SD: Standard deviation. \*\* $p < .01$ ° $p < .1$

### 4.4. Delivery

No significant main effect or interaction emerged neither for RewP or FRN, but the AD group tended to show a less pronounced RewP at the selected fronto-central electrodes for the monetary condition (p=.065, d=.-45). For details see Figure 3 and Table 2.

**Figure 2.**
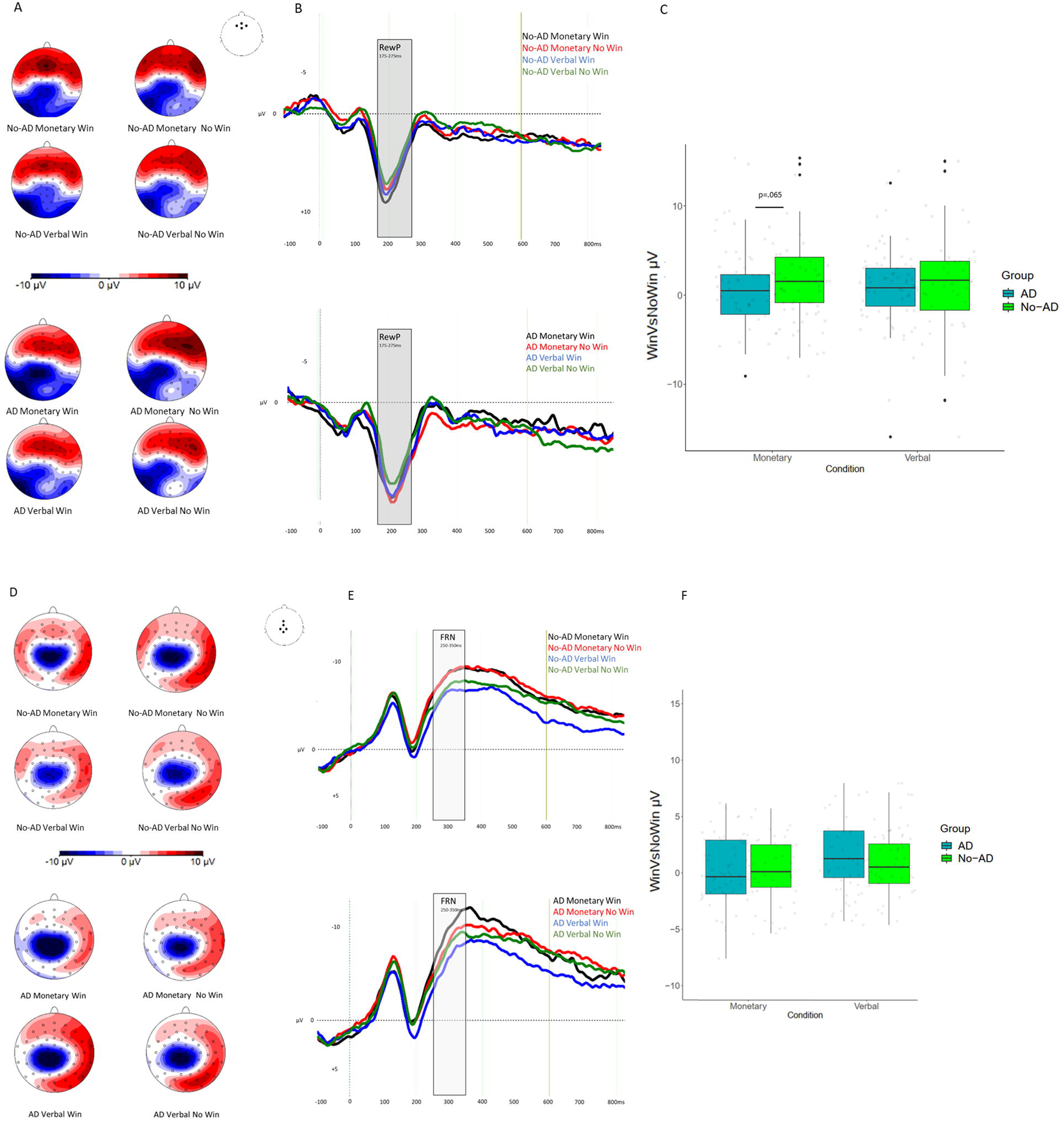
Reward Delivery Reward positivity 175-275 ms. A) Displays the topographies for No-AD (Top) and AD (bottom) for each condition. B) Displays the ERPs for No-AD (Top) and AD (bottom) for each condition. C) Extracted difference between Win and No win for each condition by group. Feedback related negativity (FRN) 250 – 350 ms. D) Displays the topographies for No-AD (Top) and AD (bottom) for each condition. E) Displays the ERPs for No-AD (Top) and AD (bottom) for each condition. F) Extracted difference between Win and No win for each condition by group. AD=Affective dysregulation

**Figure 3.**
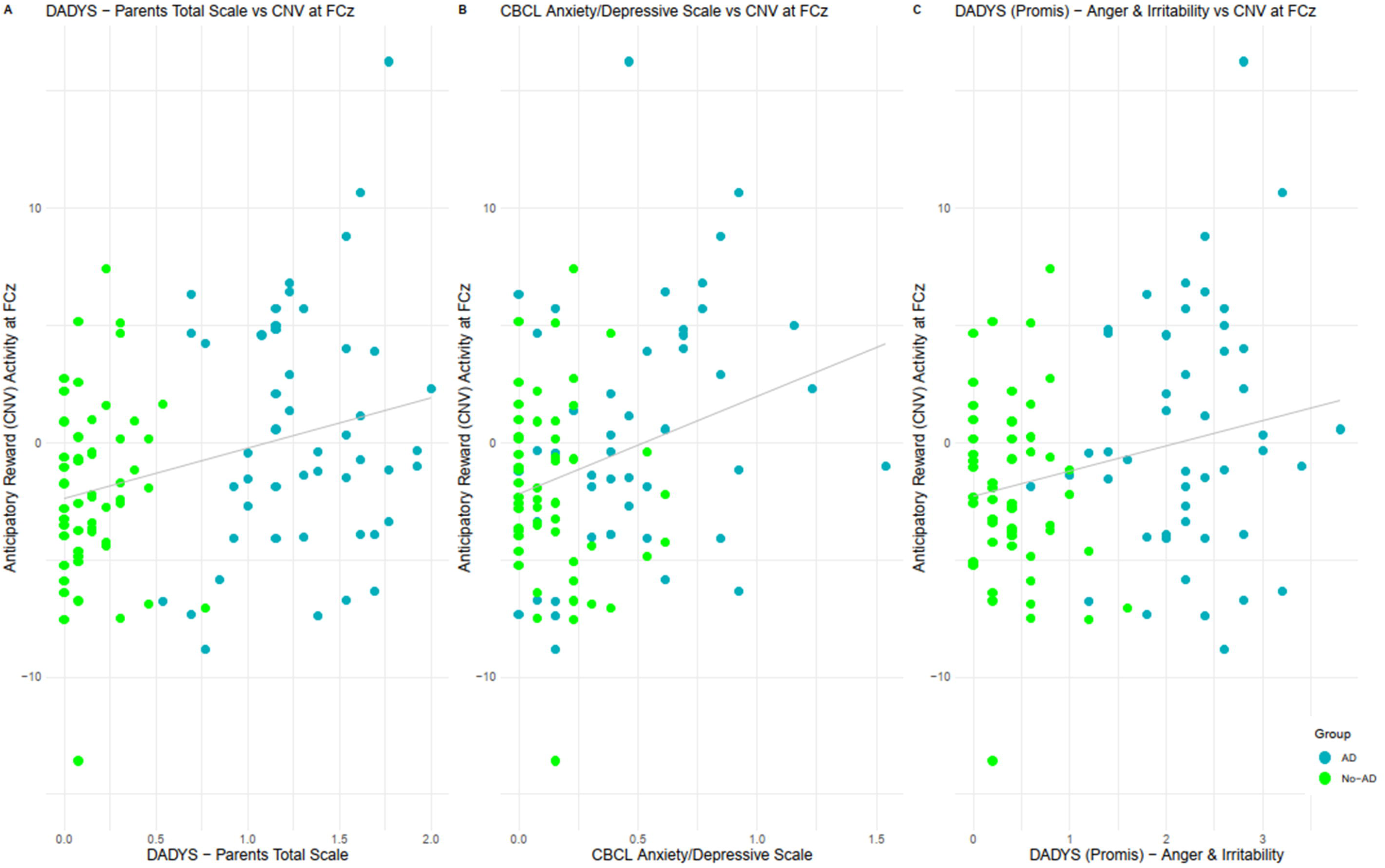
Scatter plot – Regression analyses A) Scatter plot showing a positive association between AD core symptoms and the anticipatory reward (CNV) activity at FCz (b = 0.251, p = 0.0105). B) Showing a positive association between the anticipatory reward (CNV) activity at FCz and Anxiety/Depression symptoms (b = 0.301, p = 0.003) measured via the CBCL. C) Showing a positive association between the anticipatory reward (CNV) activity at FCz and Anger and Irritability symptoms (b = 0.299, p = 0.004). AD=Affective dysregulation.

### 4.5. Sensitivity analysis

We additionally performed a sensitivity analysis, including medication and site as a separate covariate and using only participants with available behavioral data, which all did not impact the above mentioned results.

### 4.6. Regression analysis

In the first step, including only sex and medication to predict AD symptoms, the model was significant [R^2^ = 0.116, F(2,92)=6.029, p = 0.003], explaining 11% of the variance. Only medication was a significant predictor of AD symptoms (b = 0.251, p = 0.0105). In the second step, incorporating also the anticipatory reward CNV from FCz and Cz, resulted in a 7.8% improvement in the model [R^2^ change = 0.078, F(2,90)=4,375, p = 0.015]. Only the CNV activity for the monetary condition at FCz was a significant predictor of AD symptoms (b = 0.283, t=2.728 p = 0.008, Figure 3 A). In the final step, we included the delivery phase ERP activity, which did not significantly improve the model (p = 0.161). When repeating the regression analysis, but including ADHD, aggression and anxiety/depression symptoms, in a second step, 79% of the variance was explained. Including the anticipatory reward CNV from FCz and Cz increased the model significantly by 1.7% (R^2^ change = 0.017, p=0.023). The delivery phase ERPs yielded no significant model improvement (p>0.525). For details, see supplementary Table 1 & 2.

In a further stepwise regression model, we examined the dimensions related to the significant diminished reward anticipation activity at FCz. We observed that anxiety and depressive symptoms (b = 0.301, t = 3.049, p = 0.003, Figure 3 B) and anger and irritability (b = 0.299, t = 2.947, p = 0.004, Figure 3 C), accounting for 10% and 9,3% of the variance, were significantly associated with FCz activity. These associations also withstood correction for multiple testing. Regarding RewP and FRN, we found a significant negative association also with anxiety and depressive symptoms (b = -0.259, t = -2.581, p = 0.011) and RewP, although this did not withstand correction for multiple testing and should be interpreted as explorative. For details, see supplementary Table 3.

## 5. Discussion

The current study examined differences in anticipatory and delivery-related reward processing between children with and without AD, and investigated their associations with well-characterized AD symptoms. This provides a deeper understanding of the conceptualization of AD and its neurobiological mechanisms.

Concerning reward anticipation processes, this study revealed a reduced anticipatory CNV amplitude in AD, specifically at fronto-central electrodes. The reduction was most noticeable in the monetary cue condition, indicating a compromised or less active anticipatory reward process in AD children, suggesting compromised processing in the positive valence domain of the RDoC framework. Deficiencies in reward anticipation potentially reflect a blunted attribution of expectancy and motivational salience (Hägele et al., 2015) for rewarding stimuli which might contribute to difficulties in reward learning processes (Adleman et al., 2011). Furthermore, as previously shown, children with AD showed impaired ability to process conflicting emotional information (Giller et al., 2022). This impairment might extend to the processing of reward-related cues resulting in difficulties selecting an appropriate response and partially mediating also frustrative non-reward responses (Brotman et al., 2017), and contribute to unrealistic expectations, which might cause temper outburst when the expected reward fails to materialize (Wakschlag et al., 2012).

Moreover, our reward-delivery findings revealed no significant differences in RewP or FRN between the AD and No-AD. The lack of significant differences might be due to the high heterogeneity across studies and designs investigating the RewP or FRN (i.e. Hennefield et al., 2022; Proudfit, 2015; Zubovics et al., 2021). Nonetheless, the trend-level difference indicating slightly less pronounced RewP in the AD group for the monetary condition should be investigated in future larger studies, as a reduced RewP might also imply impaired reward learning (Adleman et al., 2011) and reward sensitivity (Perlman et al., 2015) consistent with the frustrative non-reward hypothesis.

In line with a dimensional perspective (i.e., Hierarchical Taxonomy of Psychopathology and RDoC), classified as a high-priority research goal (Leibenluft et al., 2024), we performed several regression analyses. First, we showed that the deviant CNV reward anticipation activity explained an additional 7.8% of the variance in AD after including medication and sex but only an additional 1.7% after including core AD symptoms, informing the limited additional diagnostic utility of the EEG assessment. However, anger/irritability and anxiety and depressive symptoms predicted a reduced CNV reward anticipation activity at FCz. This suggests that these symptoms contribute to deficits in reward expectation and anticipation, being in line with the studies linking reduced anticipatory activity with anhedonia and depressive symptoms (Ait Oumeziane et al., 2019; Stringaris et al., 2015). Furthermore, the prevailing consensus suggests that youth irritability predicts internalizing disorders in later life, particularly depression (for a meta-analysis, see Vidal-Ribas et al., 2016). It has also been proposed that irritability is a mood manifestation that shares common risk factors with depressive and anxiety disorders (Stringaris & Taylor, 2015). Interestingly, in our study, only reward anticipation demonstrated a robust link with internalizing symptoms. In contrast to prior research findings (i.e. Belden et al., 2016; Bress et al., 2013; Burkhouse et al., 2017), the RewP showed a weaker association. Our results highlight that anger/irritability and depressive and anxiety symptoms are linked to blunted reward anticipation and expectation. Since AD is a highly transdiagnostic concept associated with both externalizing and internalizing symptoms (Leibenluft et al., 2024), deficits in reward processing at a neural level might point to an important contribution of internalizing problems, probably reflecting tonic irritability (Sorcher et al., 2024). This insight may not only enable us to improve and tailor diagnostics and develop more targeted therapies, such as by systematically increasing engagement in rewarding social and interpersonal activities (Solomonov, 2023) and/or with a treatment targeting reward anticipation and motivation through pleasurable activities and imagining positive future outcomes (Craske et al., 2023), it might also suggest that a diminished neural response to anticipated reward could serve as an early indicator for developing depressive and anxiety disorders in later life and possibly inform early prevention and personalized treatment strategies.

However, this research is not without limitations. The high rate of males in the AD group might point to potential gender bias, which warrants further exploration. The RewP and FRN are commonly used in an exchangeable manner within a heterogeneous time-range (i.e. Hennefield et al., 2022; Proudfit, 2015; Zubovics et al., 2021), which we addressed by covering both. Furthermore, the reward anticipation task used was not designed to test frustration per se. The link to frustrative nonreward should be followed up with specific frustration tasks in the future, such as the affective Posner cueing task. Lastly, longitudinal data would be needed to clarify how the relation between reward anticipation, AD and internalizing symptom evolves over time.

In conclusion, our findings contribute to the body of evidence on altered reward processing in AD by implicating deviant neural activity in anticipation of reward preceding processes related to the delivery of the reward itself. Furthermore, our results underscore the importance of dimensional analyses which showed that both anger/irritability and anxiety/depressive symptoms are relevant dimensions in the pathophysiology of AD for atypical reward anticipation. This understanding offers promising directions for future research and might improve diagnostics and future interventions.

## Notes

### Competing Interest Statement

The authors have declared no competing interest.

