## Supplement for "Reward processing in children with Affective Dysregulation"

Supplementary Table 1 - Regression analysis

To which extent EEG reward-related activity can explain AD symptoms

| Coefficientsa | | | | | | | | |
| --- | --- | --- | --- | --- | --- | --- | --- | --- |
| Model | | Unstandardized Coefficients | | Standardized Coefficients | t | Sig. | Collinearity Statistics | |
|  |  | B | Std. Error | Beta |  |  | Tolerance | VIF |
| 1 | (Constant) | .962 | .201 |  | 4.787 | .000 |  |  |
|  | Sex | -.222 | .127 | -.176 | -1.745 | .084 | .941 | 1.063 |
|  | Medication | .513 | .206 | .251 | 2.488 | .015 | .941 | 1.063 |
| 2 | (Constant) | 1.026 | .195 |  | 5.247 | .000 |  |  |
|  | Sex | -.233 | .124 | -.185 | -1.882 | .063 | .928 | 1.077 |
|  | Medication | .554 | .200 | .271 | 2.773 | .007 | .936 | 1.069 |
|  | Cz_CNV_Cue Monetary | -.001 | .014 | -.006 | -.061 | .952 | .827 | 1.209 |
|  | FCZ_CNV_Cue Monetary | .042 | .016 | .283 | 2.728 | .**008** | .830 | 1.205 |
| 3 | (Constant) | 1.008 | .194 |  | 5.184 | .000 |  |  |
|  | Sex | -.227 | .125 | -.180 | -1.822 | .072 | .895 | 1.118 |
|  | Med | .554 | .199 | .271 | 2.785 | .007 | .926 | 1.080 |
|  | Cz_CNV_Cue Monetary | -.007 | .014 | -.050 | -.473 | .637 | .786 | 1.272 |
|  | FCZ_CNV_Cue Monetary | .040 | .016 | .270 | 2.560 | .**012** | .791 | 1.264 |
|  | RewP_Monetary | -.028 | .015 | -.187 | -1.815 | .073 | .830 | 1.205 |
|  | RewP_Verbal | .015 | .014 | .115 | 1.136 | .259 | .857 | 1.167 |
| a. Dependent Variable: DADYS – Parents total scale | | | | | | | | |

Model 1: F(2,94)=6.029, p=.003

Model 2: F(4,94)=5.423, p=.001

Model 3: F(6,94)=4.306, p=.001

Supplementary Table 2

Additional regression analysis including ADHD, Aggression and Anxiety/Depressive symptoms

| Model Summarye | | | | | | | | | |
| --- | --- | --- | --- | --- | --- | --- | --- | --- | --- |
| Model | R | R Square | Adjusted R Square | Std. Error of the Estimate | Change Statistics | | | | |
|  |  |  |  |  | R Square Change | F Change | df1 | df2 | Sig. F Change |
| 1 | .340a | 0.116 | 0.097 | 0.5986750 | 0.116 | 6.029 | 2 | 92 | 0.003 |
| 2 | .890b | 0.793 | 0.781 | 0.2945494 | 0.677 | 97.020 | 3 | 89 | 0.000 |
| 3 | .900c | 0.810 | 0.795 | 0.2852885 | 0.017 | 3.936 | 2 | 87 | 0.023 |
| 4 | .902d | 0.813 | 0.793 | 0.2864472 | 0.003 | 0.649 | 2 | 85 | 0.525 |
| a. Predictors: (Constant), Medication, Sex | | | | | | | | | |
| b. Predictors: (Constant), Medication, Sex, CBCL Anxiety/Depressive - T1, FBB-ADHD Total scale ADHS - T1, CBCL Aggressive behavior - T1 | | | | | | | | | |
| c. Predictors: (Constant), Medication, Sex, CBCL Anxiety/Depressive - T1, FBB-ADHS Total scale - T1, CBCL Aggressive behavior - T1, Cz_CNV_CueWin, FCZ_CNV_CueWin | | | | | | | | | |
| d. Predictors: (Constant), Medication, Sex, CBCL Anxiety/Depressive - T1, FBB-ADHS Total scale - T1, CBCL Aggressive behavior - T1, Cz_CNV_CueWin, FCZ_CNV_CueWin, RewP_Verbal, RewP_Monetary | | | | | | | | | |
| e. Dependent Variable: DADYS-Parent Total scale - T1 | | | | | | | | | |

| Coefficientsa | | | | | | | | |
| --- | --- | --- | --- | --- | --- | --- | --- | --- |
| Model | | Unstandardized Coefficients | | Standardized Coefficients | t | Sig. | Collinearity Statistics | |
|  |  | B | Std. Error | Beta |  |  | Tolerance | VIF |
| 1 | (Constant) | 0.962 | 0.201 |  | 4.787 | 0.000 |  |  |
|  | Sex | -0.222 | 0.127 | -0.176 | -1.745 | 0.084 | 0.941 | 1.063 |
|  | Medication | 0.513 | 0.206 | 0.251 | 2.488 | 0.015 | 0.941 | 1.063 |
| 2 | (Constant) | 0.136 | 0.113 |  | 1.207 | 0.231 |  |  |
|  | Sex | -0.015 | 0.065 | -0.012 | -0.232 | 0.817 | 0.865 | 1.157 |
|  | Medication | 0.189 | 0.106 | 0.093 | 1.794 | 0.076 | 0.869 | 1.150 |
|  | FBB-ADHS Total scale - T1 | 0.030 | 0.068 | 0.032 | 0.442 | 0.659 | 0.453 | 2.207 |
|  | CBCL Aggressive behavior- T1 | 1.054 | 0.106 | 0.739 | 9.895 | 0.000 | 0.417 | 2.396 |
|  | CBCL Anxiety/Depressive- T1 | 0.315 | 0.118 | 0.160 | 2.676 | 0.009 | 0.650 | 1.539 |
| 3 | (Constant) | 0.186 | 0.111 |  | 1.682 | 0.096 |  |  |
|  | Sex | -0.019 | 0.064 | -0.015 | -0.296 | 0.768 | 0.855 | 1.169 |
|  | Medication | 0.214 | 0.103 | 0.105 | 2.088 | 0.040 | 0.863 | 1.159 |
|  | FBB-ADHS Total scale - T1 | 0.031 | 0.066 | 0.033 | 0.471 | 0.639 | 0.451 | 2.219 |
|  | CBCL Aggressive behavior- T1 | 1.055 | 0.104 | 0.739 | 10.186 | 0.000 | 0.414 | 2.413 |
|  | CBCL Anxiety/Depressive- T1 | 0.240 | 0.118 | 0.122 | 2.039 | 0.044 | 0.612 | 1.634 |
|  | Cz_CNV_Cue Monetary | 0.000 | 0.007 | 0.001 | 0.023 | 0.982 | 0.817 | 1.225 |
|  | FCZ_CNV_Cue Monetary | 0.020 | 0.008 | 0.136 | 2.550 | **0.013** | 0.764 | 1.309 |
| 4 | (Constant) | 0.196 | 0.112 |  | 1.755 | 0.083 |  |  |
|  | Sex | -0.026 | 0.065 | -0.021 | -0.398 | 0.692 | 0.821 | 1.219 |
|  | Medication | 0.225 | 0.104 | 0.110 | 2.168 | 0.033 | 0.848 | 1.179 |
|  | FBB-ADHS Total scale - T1 | 0.023 | 0.068 | 0.024 | 0.337 | 0.737 | 0.438 | 2.285 |
|  | CBCL Aggressive behavior- T1 | 1.060 | 0.104 | 0.743 | 10.145 | 0.000 | 0.410 | 2.436 |
|  | CBCL Anxiety/Depressive- T1 | 0.226 | 0.120 | 0.114 | 1.877 | 0.064 | 0.591 | 1.691 |
|  | Cz_CNV_Cue Monetary | -0.002 | 0.007 | -0.013 | -0.247 | 0.806 | 0.768 | 1.302 |
|  | FCZ_CNV_Cue Monetary | 0.021 | 0.008 | 0.142 | 2.605 | **0.011** | 0.736 | 1.358 |
|  | RewP_Monetary | -0.006 | 0.008 | -0.038 | -0.707 | 0.481 | 0.783 | 1.278 |
|  | RewP_Verbal | 0.007 | 0.007 | 0.054 | 1.056 | 0.294 | 0.830 | 1.204 |
| a. Dependent Variable: DADYS-Parent Total scale - T1 | | | | | | | | |

Supplementary Table 3 - Predicting CNV activity

| Predicting CNV activity – Regression analyses | | | | |
| --- | --- | --- | --- | --- |
|  | Reward anticipation CNV at FCz | | |  |
| Predictors | beta | t | p | Model p |
| DADYS Irritability/Impulsivity | .249 | 2.360 | .020 | .096 |
| DADYS Promis (Anger/Irritability) | .299 | 2.947 | **.004** | **.026** |
| DADYS Exhuberance | .238 | 2.317 | .023 | .095 |
| DADYS Function | .246 | 2.254 | .027 | .114 |
| DADYS Positive emotionality | -102 | -.925 | .357 | .624 |
| CBCL Aggression | .194 | 1.799 | .075 | .250 |
| FBB ADHD | .241 | 2.267 | .026 | .115 |
| CBCL Anxiety/Depression | .301 | 3.064 | **.003** | **.019** |
|  | Reward delivery RewP | |  |  |
| DADYS Irritability/Impulsivity | -.136 | -1.242 | .218 | .298 |
| DADYS Promis (Anger/Irritability) | -.147 | -1.379 | .171 | .258 |
| DADYS Exhuberance | -.204 | -1.950 | .054 | .153 |
| DADYS Function | -.231 | -2.062 | .042 | 0.96 |
| DADYS Positive emotionality | .120 | 1.077 | .284 | .322 |
| DADYS Total score | -.134 | -1.234 | .221 | .301 |
| CBCL Aggression | -.182 | -1.688 | .095 | .175 |
| FBB ADHD | -.086 | -.775 | .440 | .433 |
| CBCL Anxiety/Depression | -.259 | -2.581 | .011 | .035 |
|  | Reward delivery FRN | |  |  |
| DADYS Irritability/Impulsivity | .001 | .012 | .990 | .571 |
| DADYS Promis (Anger/Irritability) | -.045 | -.419 | .676 | .535 |
| DADYS Exhuberance | -.079 | -.750 | .455 | .440 |
| DADYS Function | .002 | .016 | .988 | .495 |
| DADYS Positive emotionality | -.015 | -.134 | .893 | .510 |
| DADYS Total score | -.003 | -.026 | .979 | .571 |
| CBCL Aggression | -.001 | -.010 | .992 | .571 |
| FBB ADHD | -002 | -019 | -985 | -571 |
| CBCL Anxiety/Depression | .027 | .260 | .796 | .557 |

In Bold: CBCL Anxiety/Depression scale and the DADYS Promis (Anger/Irritability)

significantly predicted (after correction for multiple testing) reward anticipation deficits measured at FCz.

Supplementary Figure 1 – Reward anticipation task

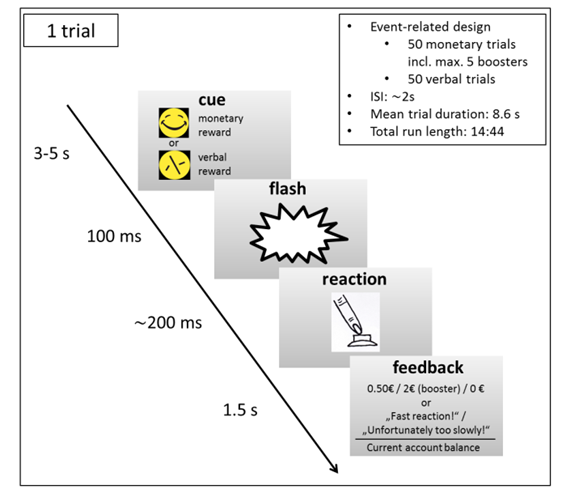

The reward anticipation task requires a fast button press to a flash cued by a smiley or a scrambled smiley indicating monetary or verbal feedback.

Supplementary Figure 2 - Reaction time

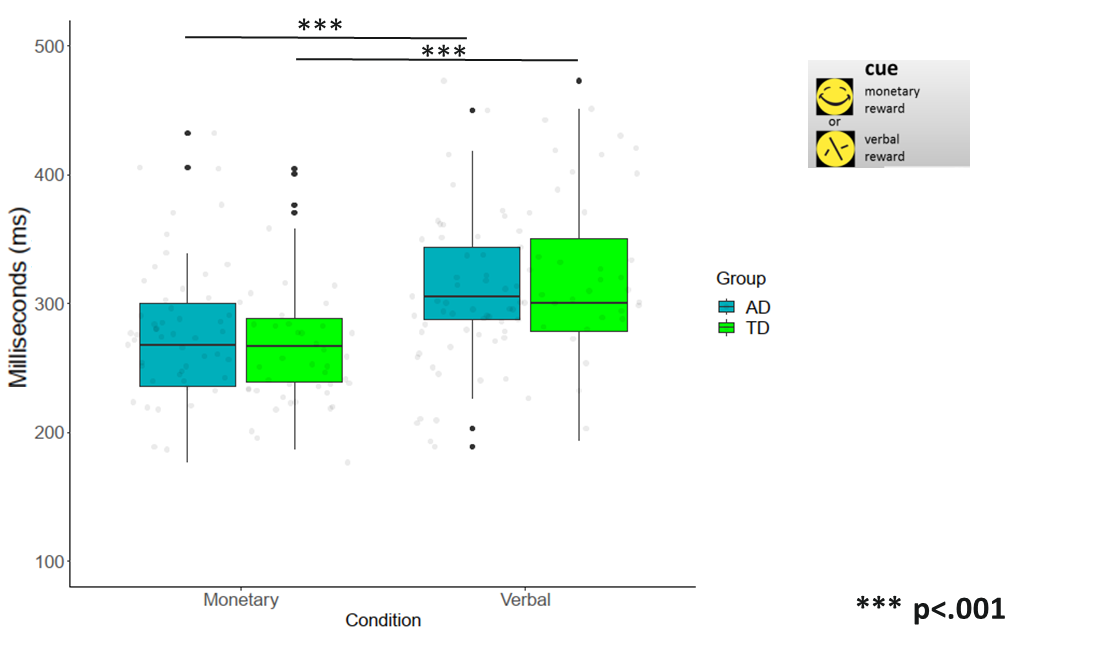
